# Islands of Signal and Transcriptomic Sequencing: A Foundation Model for Mutation and Lineage Prediction based on DNA Methylation and RNA-seq

**DOI:** 10.64898/2025.12.01.691534

**Authors:** Alexandros Alexakos, Aristotelis Tsirigos

## Abstract

DNA methylation and RNA-seq provide complementary views of oncogenic state, but their high dimensionality complicates robust modeling. We develop a pancancer, multiomic foundation model that jointly encodes CpG-island DNA methylation and gene expression from TCGA, TARGET, CPTAC-3, and HCMI. Probe-level methylation is aggregated into CpG-island features, and RNA-seq is reduced to high-variance genes, yielding compact inputs for modality-specific MLP encoders. A BERT-like transformer with masked reconstruction and cross-modal prediction objectives learns a shared embedding space that supports missing-modality inputs.

We evaluate the learned representations in two zero-shot settings: (i) cancer-type classification using a linear probe on frozen embeddings, and (ii) mutation prediction for 214 genes using a shallow MLP. The model achieves high performance for many tumor types and gene-cancer pairs without encoder finetuning. Pathway-level analyses show that hallmark oncogenic and immune programs appear as smooth gradients in the embedding space, indicating that the model captures biologically meaningful structure. These results demonstrate that combining CpG-island grouping with multiomic foundation pre-training yields compact, informative embeddings for mutation and lineage inference across cancers.

## 1 Introduction

DNA methylation is a central epigenetic mechanism that regulates gene expression and maintains genomic stability. Unusual methylation patterns are frequently observed across cancers and have been linked to oncogene activation, tumor suppressor silencing, and global genomic instability [1, 2]. Because methylation changes are both widespread and tumor-type specific, methylation data have emerged as a powerful omic for biomarker discovery and predictive modeling [3]. In parallel, bulk RNA sequencing has become the standard approach for characterizing tumor transcriptomes, providing a complementary readout of pathway activation, cell-state composition, and oncogenic signaling [4, 5].

One promising direction is the inference of somatic mutations from methylation and expression profiles [6]. While next-generation sequencing remains the gold standard for mutation calling, it is expensive, resource-intensive, and often unavailable for retrospective cohorts or routine clinical workflows [7, 8]. In contrast, array-based methylation profiling is more cost-effective and widely available, and paired RNA sequencing is increasingly common across large research consortia. Together, these modalities can potentially serve as surrogates for predicting the mutation status of key cancer driver genes and pathways [9]. However, two major challenges remain: (i) the high dimensionality and noise of probe-level methylation and genome-wide expression data, and (ii) poor generalization of models trained in a single cohort when applied to independent datasets, institutions, or disease subtypes.

Large public cancer genomics initiatives now provide an opportunity to address these issues at scale. Pan-cancer efforts such as The Cancer Genome Atlas (TCGA), the Clinical Proteomic Tumor Analysis Consortium (CPTAC-3), the Human Cancer Models Initiative (HCMI), and the Therapeutically Applicable Research to Generate Effective Treatments program (TARGET) collectively offer thousands of samples with harmonized somatic mutation calls, DNA methylation, and RNA-seq profiles across adult and pediatric tumor types [10, 11, 12, 13]. At the same time, the machine learning community has converged on the paradigm of *foundation models*: large models pre-trained on broad data that can be adapted to many downstream tasks through zero-shot evaluation or lightweight fine-tuning [14, 15]. Bringing these ideas together suggests a natural strategy for cancer genomics: train a pan-cancer, multi-omic foundation model once on all available public datasets, then reuse it to solve diverse mutation-prediction tasks with minimal task-specific supervision.

To address the challenges of dimensionality and generalization in methylation-based mutation prediction, we introduce a framework centered on three methodological advances:

1. **CpG-to-island grouping.** Instead of using hundreds of thousands of probe-level CpGs, we aggregate them into biologically meaningful CpG islands. This reduces dimensionality by an order of magnitude while preserving regulatory signal, mitigating platform-specific noise and overfitting risks, and yielding a stable feature space that can be shared across datasets and cancer types.
2. **Pan-cancer multiomic foundation model pre-training.** We construct a pan-cancer encoder jointly trained on CpG-island methylation and matched RNA-seq expression from all available public cohorts (TCGA, CPTAC-3, HCMI, TARGET). The encoder is optimized to learn representations that are predictive of somatic mutations across many genes and tumor types, following the foundation-model paradigm of broad pre-training followed by task adaptation [14, 15, 10, 11].
3. **Zero-shot evaluation and cancer-specific fine-tuning.** We evaluate the pre-trained encoder in two regimes. First, we perform zero-shot mutation prediction using simple downstream heads (e.g., *k*-nearest neighbors and logistic regression) trained on the learned representations without updating the encoder. Second, we fine-tune the encoder and a logistic head on cancer-specific subsets and evaluate on independent validation cohorts, comparing performance against stand-alone deep learning models trained from scratch on each cancer type.

**Our contributions** are threefold: (i) we demonstrate that CpG-island grouping and rna-seq genes reduction with column variation filtering reduces dimensionality by ∼90% while maintaining predictive information for mutation status and cancer type prediction; (ii) we establish a foundation model based on CpG islands beta values and RNA-seq TPM values that is able to predict driver mutations and cancer types; (iii) we evaluate these findings through pathway enrichment to state the biological insight except from the technological. Together, these findings highlight the potential of dimensionality-aware feature engineering and foundation-model transfer learning to enable robust, generalizable mutation prediction from DNA methylation and RNA-seq across cancers.

## 2 Background

Predicting cancer mutations through DNA methylation offers a cost-effective alternative to sequencing-based approaches [16, 17]. Recent studies demonstrate that methylation patterns can predict driver mutations with high accuracy across multiple cancer types. For instance, *IDH1* - mutant gliomas exhibit the distinctive G-CIMP phenotype detectable through methylation arrays [18], while pan-cancer analyses reveal that methylation profiles predict recurrent alterations in *TP53*, *BRAF*, and other driver genes nearly as effectively as gene expression data [19, 20]. The relationship between methylation and mutation status appears bidirectional, with mutations influencing methylation patterns and methylation changes potentially predisposing to specific mutational processes [21].

In parallel, bulk RNA sequencing has become the standard approach for characterizing tumor transcriptomes, providing a complementary readout of pathway activation, cell-state composition, and oncogenic signaling [4, 5, 22, 23]. Pan-cancer comparisons of multiple omics layers have shown that gene expression and DNA methylation are consistently among the most informative modalities for predicting somatic mutation states across a broad panel of cancer genes, often outperforming other functional readouts [19]. These observations motivate multiomic models that jointly exploit methylation and RNA-seq to infer mutation status, especially in settings where direct sequencing is unavailable or incomplete.

The primary challenge in methylation-based prediction lies in the extreme dimensionality of array data, with platforms measuring 450,000–850,000 CpG sites per sample. Various dimensionality reduction strategies have been proposed, with CpG island (CGI) grouping emerging as particularly effective. By aggregating probes within biologically meaningful CGIs, studies achieve 90–95% feature reduction while preserving regulatory signals [24, 25]. This approach aligns with biological understanding, as CGIs represent functional regulatory units where methylation changes often occur coordinately [26, 27]. Advanced methods like beta-mixture models collapse thousands of CpGs into interpretable parameters [24], while network-guided approaches leverage co-methylation patterns and graph structure to define modules of coordinated methylation [28]. Recently, MethylCapsNet and related capsule-based architectures have demonstrated that explicit CGI or region-level grouping enhances both interpretability and predictive performance for methylation-based classifiers [29, 30].

Transfer learning and foundation models are becoming increasingly central in genomics and methylation analysis. Genomic foundation models trained on massive datasets demonstrate that pre-training captures generalizable patterns transferable across tasks, tissues, and disease contexts [31, 14, 15, 32]. In methylation specifically, approaches such as *cfMethylPre* have reported superior liquid biopsy performance by pre-training on dozens of cancer types [33], while MethylGPT and CpGPT—transformer architectures trained on large collections of methylomes—illustrate that sequence- and region-aware models can capture complex dependencies in methylation profiles [34, 35]. More broadly, deep learning and transfer learning have been leveraged for multiomics integration, combining methylation, expression, and additional data types to improve prediction of survival, subtypes, and treatment response [36].

Multi-modal approaches integrating methylation with other omics data further improve prediction accuracy and biological interpretability. Capper et al. established a comprehensive methylation-based classification system for central nervous system tumors, demonstrating clinical utility and robustness across institutions [37]. MethNet extends this paradigm by integrating large-scale methylation and gene expression data across multiple cancers to identify distal regulatory hubs that influence transcription and patient outcome [38]. These studies highlight the value of combining methylation with transcriptomic readouts to capture regulatory programs downstream of genetic alterations.

Despite these advances, no prior work has systematically integrated CpG island grouping, pan-cancer multiomic (methylation + RNA-seq) pre-training strategies specifically for mutation prediction. Existing TCGA-wide studies of mutation prediction typically rely on probe-level methylation features without biological grouping, or treat methylation and expression independently rather than through a unified multi-modal encoder [19]. Similarly, methylation-focused foundation models have largely targeted imputation, age prediction, or cell-type deconvolution, rather than direct somatic mutation inference [34, 35, 33]. Our study addresses this gap by combining biologically informed CGI aggregation with TCGA-wide, pan-cancer foundation model pre-training on both DNA methylation and RNA-seq, followed by cancer-specific fine-tuning and zero-shot evaluation on external cohorts. This approach leverages both the dimensionality reduction benefits of CGI grouping and the generalization power of foundation-model transfer learning to achieve robust mutation prediction across diverse cancer types and independent datasets.

## 3 Results

In this section, we present the results of the CpG islands’ grouping reduction rate, their individual per cancer training models and their external validation, and finally their comparison to the foundation model prediction for the same external validation datasets.

### 3.1 Preprocess and Methodoly

#### 3.1.1 Dimensionality Reduction

To ensure that both modalities could be modeled efficiently within the multi-modal foundation architecture, we applied structured dimensionality reduction pipelines to RNA-seq and DNA methylation data. These procedures preserved biologically informative signal while removing low-variance, low-coverage, or redundant features. The full pipelines are illustrated in Figures 1c, 1e, and 1d.

**Figure 1:**
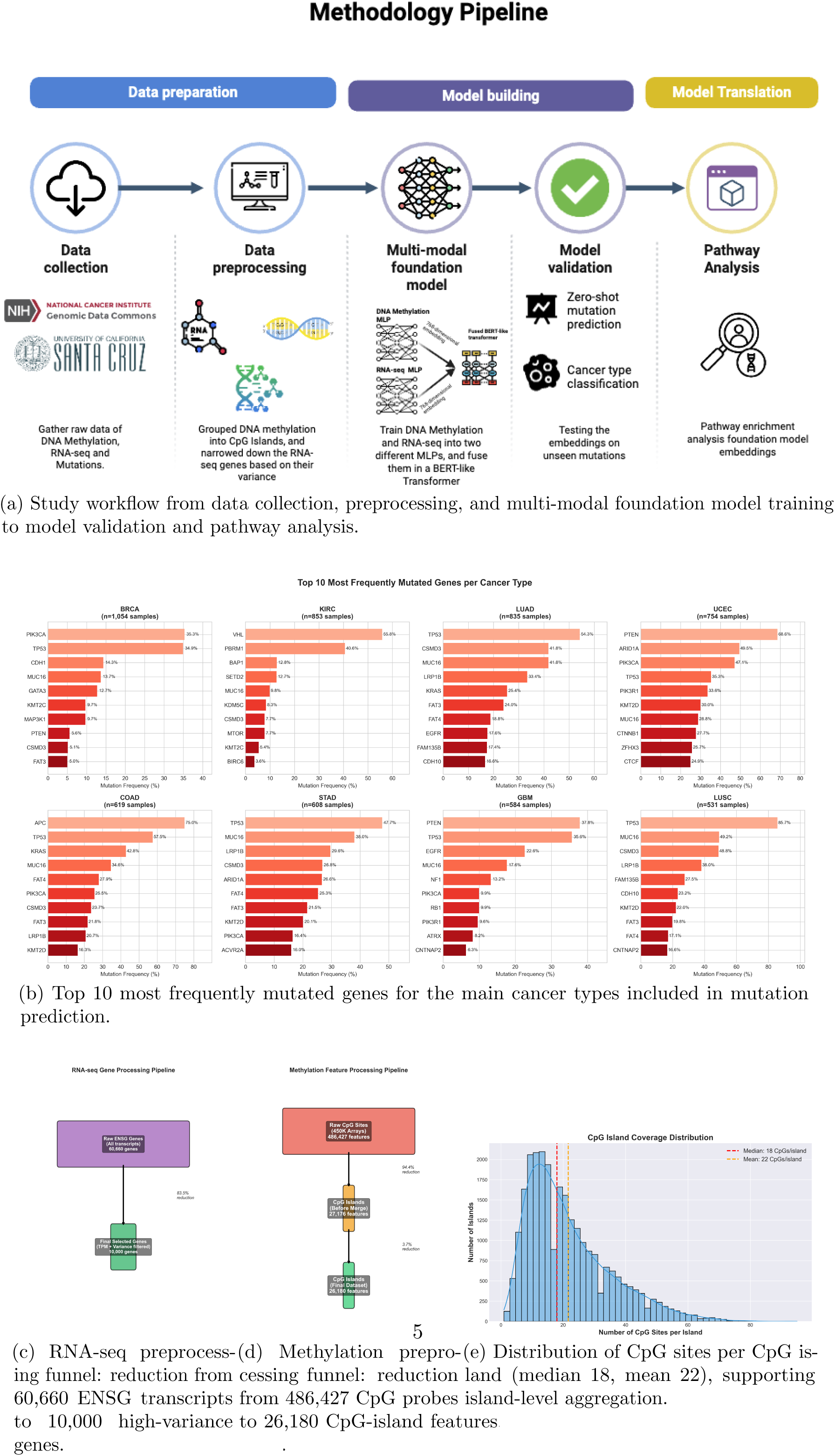
Overview of study workflow, data preprocessing, and foundation model embeddings.

Raw RNA-seq profiles obtained from GDC contain expression measurements for approximately 60,660 Ensembl transcript identifiers (ENSG). Many correspond to lowly expressed transcripts, pseudogenes, or features exhibiting negligible variation across tumors. After TPM normalization and replicate merging, we filtered genes using two criteria:

1. **Expression filtering:** removal of genes with negligible TPM values across the cohort.
2. **Variance filtering:** selection of genes exhibiting sufficient between-sample variability to contribute to downstream representation learning.

As visualized in Figure 1c, this pipeline reduced the dimensionality of the transcriptome by **83.5%**, yielding a curated set of **10,000 informative genes**. This preserves transcriptome-wide biological signal while mitigating noise and extreme sparsity.

Methylation data from Illumina 450K arrays contain ∼486,427 CpG probes, substantially exceeding the capacity of deep multi-modal architectures and containing considerable redundancy.

We therefore aggregated CpG probes into biologically meaningful **CpG islands** based on genome annotation.

Figure 1e shows that CpG islands contain a median of **18 CpG sites** (mean 22), supporting the island-level abstraction as an appropriate unit of methylation representation. Per-island beta values were computed by averaging probe-level values after quality filtering.

This produced an intermediate set of **27,176 islands**, followed by the exclusion of low-coverage islands, resulting in a final set of **26,180 CpG-island features**. As illustrated in Figure 1d, this corresponds to a **94.4% reduction** from the original probe space while retaining epigenetically relevant regulatory regions.

#### 3.1.2 Somatic Mutation Processing and Gene Selection

Somatic mutation profiles were obtained from the GDC MAF (Mutation Annotation Format) files for all TCGA projects, including all available *tumor*, *normal-adjacent*, *metastatic*, and *primary* samples. To maximize cross-cohort consistency, we used the GDC pipeline–harmonized

MAFs, which provide standardized variant calling across cancer types.

Because mutation frequency varies drastically across cancer types, we constructed a unified pan-cancer target gene list by selecting genes mutated in at least 1% of samples across the combined TCGA–TARGET–CPTAC cohorts. This threshold reduces extremely sparse labels while preserving the major driver and recurrently mutated genes.

The final gene set contains:

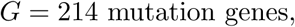

each treated as an independent binary classification task in the multi-task prediction head.

Figure 1b summarizes the top 10 most frequently mutated genes across major cancer types in our dataset, including BRCA, LUAD, LUSC, COAD, STAD, GBM, KIRC, and UCEC. These distributions highlight several key biological patterns:

- **TP53** is the dominant mutation across most cancers (e.g., 85.7% in LUSC, 54.3% in LUAD).
- **PIK3CA** is highly recurrent in breast and gynecological cancers (e.g., 35.3% in BRCA, 47.1% in UCEC).
- **VHL** and **PBRM1** define the mutational signature of KIRC (55.8% and 40.6%, respectively).
- **APC** dominates the colorectal cancer landscape (75% in COAD).
- **PTEN**, **ARID1A**, and **CTNNB1** characterize endometrial carcinomas.

These structured and cancer-specific mutational patterns underscore the importance of a multi-task framework: although each gene is treated independently for prediction, shared representation learning allows the foundation model to capture underlying co-mutation and pathway-level relationships.

The mutation matrix plays two roles in our foundation modeling pipeline:

1. As targets for supervised fine-tuning of the fused representations,
2. As labels for downstream evaluation tasks such as per-gene AUC/AP computations and cancer-type–specific performance analysis.

The multi-label structure, combined with the diverse mutation profiles across cancer types, provides a challenging and biologically rich supervision signal that tests the generalization capability of the foundation model.

### 3.2 Foundation Model

Our approach centers on a multi-modal genomic foundation model that jointly encodes DNA methylation and RNA-seq profiles using a hybrid architecture consisting of asymmetric MLP encoders and a BERT-like transformer fusion module. Each modality is first compressed into a 768-dimensional latent space through modality-specific deep encoders that account for their distinct feature scales (10,000 genes vs. 26,000 CpG islands). The resulting embeddings are fused through stacked self-attention and cross-attention layers that learn regulatory relationships within and between modalities. The model is pre-trained using masked value reconstruction and cross-modal prediction objectives, enabling it to learn robust pan-cancer representations without mutation labels. During fine-tuning, a multi-task classification head predicts mutation status for 214 genes, with support for missing modalities via learned placeholder tokens and modality-aware scaling. This design yields a flexible, biologically informed foundation model capable of generalizing across cancer types, platforms, and incomplete data settings.

### 3.3 UMAP Visualization of the Multi-Modal Embedding Space

To assess the structure learned by the foundation model and evaluate whether the fused representations capture meaningful biological variation, we projected the final joint embeddings into two dimensions using Uniform Manifold Approximation and Projection (UMAP). The resulting two-panel visualization is shown in Figure 2a.

**Figure 2:**
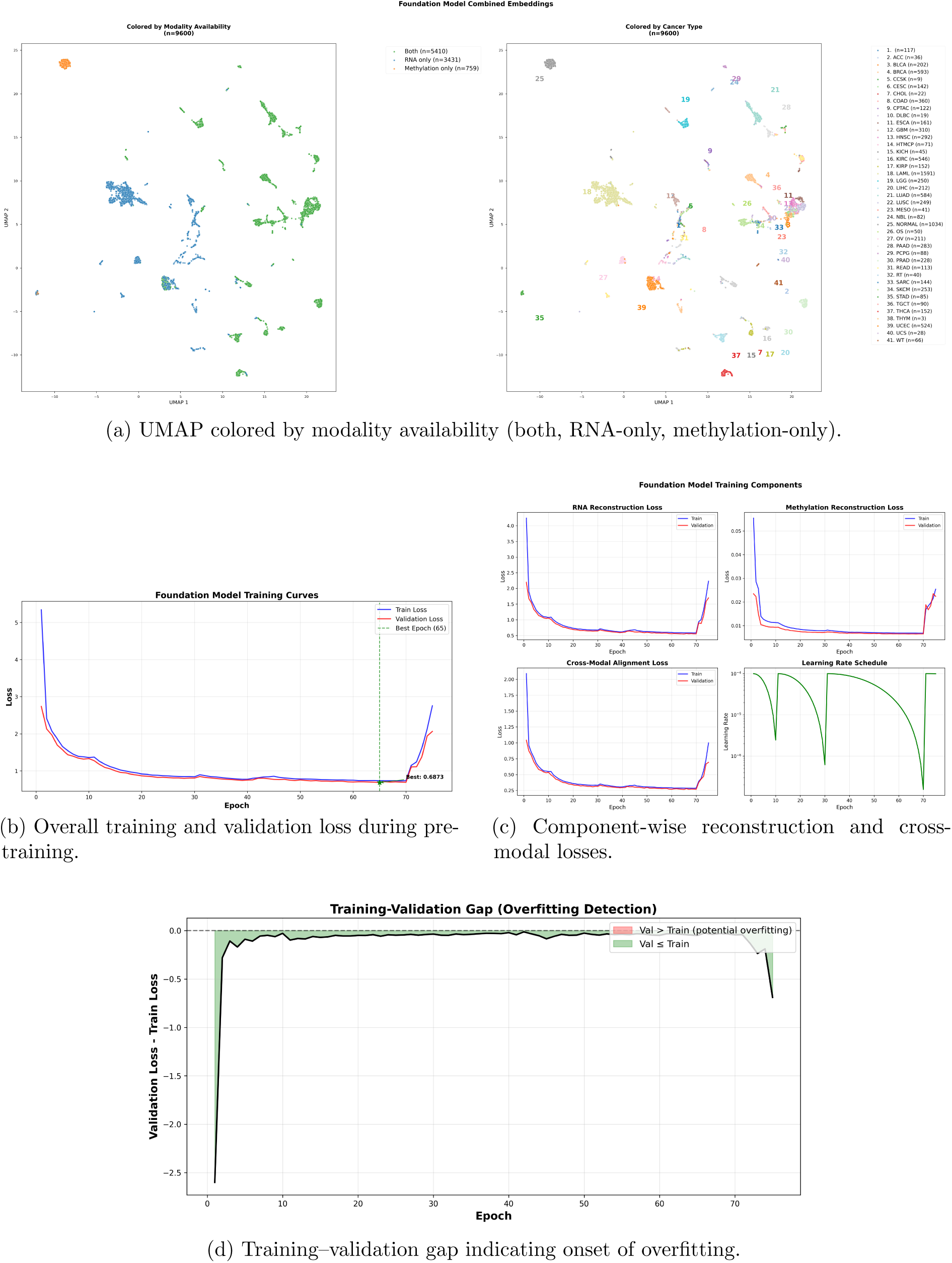
Training dynamics of the multi-modal foundation model. Top-left: overall training and validation loss curves with early-stopping epoch highlighted. Top-right and bottom-left: component-wise losses for RNA reconstruction, methylation reconstruction, and cross-modal prediction. Bottom-right: train–validation gap used for overfitting detection.

In the first panel of Figure 2a, samples are colored according to modality availability: both RNA-seq and methylation, RNA-only, or methylation-only. Samples with both modalities form the most compact and coherent manifold, reflecting the richer information content of fully paired profiles. RNA-only samples occupy a broader submanifold, consistent with higher variance in transcriptomic profiles, whereas methylation-only samples cluster more tightly, reflecting the relative stability of CpG-island methylation. Importantly, all three configurations remain aligned within a shared latent space, demonstrating that the learned missing-modality tokens and modality-aware scaling enable robust embedding of incomplete inputs.

The second panel of Figure 2a colors the same embeddings by cancer type. Tumors form clear, well-separated clusters despite the fact that cancer type was *not* used during pre-training. Major groups such as gliomas (LGG, GBM), lung cancers (LUAD, LUSC), renal carcinomas (KIRC,

KIRP), and colorectal cancers (COAD, READ) appear as distinct submanifolds. Rare cancers (e.g., UVM, CHOL, ACC) remain separable despite limited sample counts. Hematological malignancies (LAML, NBL, DLBC) map to remote regions of the space, reflecting their distinct molecular signatures.

#### 3.3.1 Training Curve Analysis

The training behavior of the foundation model is summarized in Figure 2. The overall training and validation losses (Figure 2b) decrease smoothly over the course of pre-training and remain closely aligned, indicating stable optimization with minimal early overfitting. The lowest validation loss is reached at epoch 65, after which both curves rise as the cosine-annealing learning rate schedule restarts, marking the onset of overfitting and motivating early stopping at this epoch. A breakdown of the individual loss components is shown in Figure 2c. RNA reconstruction, methylation reconstruction, and cross-modal prediction losses all drop rapidly during the first epochs and then converge to a narrow plateau, demonstrating effective learning of both intra-modal structure and inter-modal relationships. RNA reconstruction remains higher in absolute magnitude, consistent with the larger dynamic range and variance of transcriptomic data, whereas methylation reconstruction converges to a lower and more stable baseline. The cross-modal loss steadily declines, confirming that the model successfully aligns RNA-seq and methylation representations during pre-training.

The training–validation gap in Figure 2d provides an additional view of generalization. After the brief burn-in period at the beginning of training, the gap stays close to zero for most epochs, indicating that validation performance closely tracks training performance. Only near the final epochs does the gap widen, coinciding with the increase in validation loss. Together, these curves show that the foundation model trains stably, achieves consistent improvements across all self-supervised objectives, and maintains good generalization up to the selected early-stopping point.

### 3.4 Evaluation

Figure 3 summarizes how well the foundation model embeddings support downstream classification tasks and how strongly they align with known biological pathways. Using a linear probe on frozen embeddings, cancer-type classification achieves high precision, recall, and F1 scores across the top 20 tumor types (Figure 3a), with many entities approaching near-perfect performance. A shallow MLP trained on top of the same embeddings in a zero-shot setting yields strong mutation prediction for selected cancer–gene pairs (Figure 3b), with several pairs (e.g. DLBC–*BCR*, ACC–*SMC1A*) reaching high F1 and ROC–AUC.

**Figure 3:**
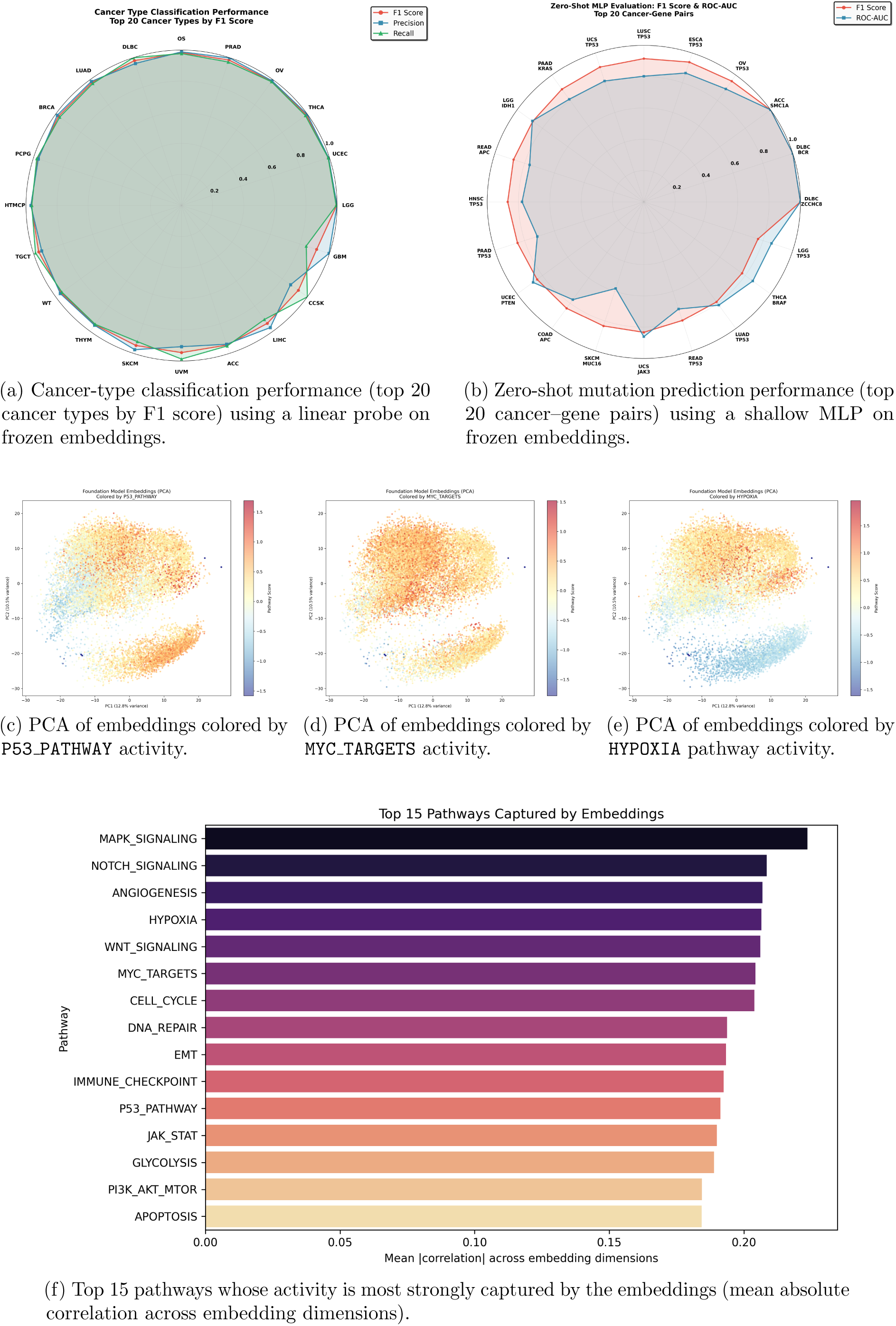
Zero-shot evaluation and pathway structure in the foundation model embeddings.

Panels 3c–3e project the joint embeddings onto the first two principal components and color samples by pathway activity for P53 PATHWAY, MYC TARGETS, and HYPOXIA. Smooth gradients across the embedding manifold indicate that major oncogenic programs vary continuously along dominant axes of variation, rather than being confined to isolated clusters. Finally, the pathway enrichment summary in Figure 3f shows that the embeddings strongly capture signaling and cancer-relevant processes such as MAPK, NOTCH, angiogenesis, hypoxia, WNT, MYC, cell cycle, DNA repair, and immune checkpoint pathways. Together, these results demonstrate that the foundation model learns embeddings that are both highly predictive for downstream tasks and tightly coupled to interpretable biological programs.

## 4 Discussion

### 4.1 Comparison to Prior Work

DNA methylation and RNA-seq have each been used independently to infer genomic alterations and to build robust cancer classifiers [1, 2, 3, 4, 5, 22, 23, 19, 37]. Prior mutation-prediction efforts, however, typically relied on probe-level methylation features alone or on single-modality, single-cohort designs, which can exacerbate overfitting and limit generalization across datasets and disease contexts [19]. Our work differs in two key ways.

First, on the methylation side, we apply biologically informed CpG-island aggregation, compressing hundreds of thousands of probes into island-level features with ∼90–95% dimensionality reduction while preserving regulatory signal. This grouping mitigates platform-specific noise and overfitting risks and yields a stable feature space that can be shared across datasets and cancer types.

Second, and critically, we treat methylation and RNA-seq as a *joint* multiomic system: CpG-island features and high-variance gene expression profiles are encoded by modality-specific MLPs and fused in a shared transformer, yielding a single representation that integrates epigenetic and transcriptomic information. This goes beyond earlier studies that typically model methylation and expression in isolation or with simple feature concatenation [19]. By pre-training this multiomic encoder on pan-cancer cohorts from TCGA, CPTAC-3, HCMI, and TARGET, we follow the broader trend of genomic foundation models that capture transferable structure across tasks, tissues, and disease contexts [31, 14, 15, 32]. In the methylation domain, approaches such as *cfMethylPre*, MethylGPT, and CpGPT have demonstrated the benefits of large-scale pre-training for downstream tasks [33, 34, 35], but these have primarily focused on methylation alone and on diagnostics, aging, or cell-type deconvolution rather than joint multiomic embeddings for mutation and lineage prediction.

### 4.2 Insights from Multi-Omic Foundation Modeling

The structure of the learned embedding space suggests that the foundation model captures coherent biological variation across both modalities. UMAP projections of the fused embeddings (Figure 2a show that samples with paired methylation and RNA-seq form a compact manifold, while RNA-only and methylation-only samples occupy consistent submanifolds that remain well-aligned within the same latent space. This behavior reflects both the richer information content of fully paired profiles and the stabilizing effect of methylation, and demonstrates that the learned missing-modality tokens successfully embed incomplete inputs without collapsing them into trivial regions of the space.

Coloring the same embeddings by cancer type reveals clear clustering by lineage, including separation of gliomas, lung cancers, renal carcinomas, colorectal cancers, and hematologic malignancies. Importantly, cancer type is never used as an explicit pre-training target; the observed clustering therefore indicates that tissue-of-origin and major molecular programs are implicitly encoded by the self-supervised objectives and the multiomic fusion of CpG islands and RNA-seq.

Zero-shot evaluations further support the quality of these embeddings for downstream tasks. A linear probe trained on frozen embeddings achieves high precision, recall, and F1 scores across the top 20 tumor types (Figure 3b, indicating that a simple classifier can recover cancer-type information from the joint representation. A shallow MLP trained on the same frozen embeddings yields strong mutation prediction for selected cancer-gene pairs (Figure 3a), with several pairs (e.g. DLBC–BCR, ACC–SMC1A) reaching high F1 and ROC AUC. Also, a notable gene that is present in the metrics of the downstream task is *TP53*, which is biologically correct since it’s one of the most mutated genes across cancers. This shows that TP53 shares same DNA methylation and RNA patterns regardless of the cancer type. Overall, the results indicate that a single multiomic foundation model can support both lineage and mutation inference without task-specific modification of the encoder, relying only on lightweight heads.

The pathway analyses reveal that the embeddings are tightly coupled to interpretable biology. PCA of the joint embeddings colored by pathway activity scores (P53, MYC targets, hypoxia; Figures 3c-3e) shows smooth gradients along dominant axes of variation, rather than isolated clusters. The pathway correlation summary (Figure 3f) highlights oncogenic and immune programs, including MAPK, NOTCH, angiogenesis, hypoxia, WNT, MYC, DNA repair, cell cycle, and immune checkpoint signaling, as among the most strongly captured. Together, these observations suggest that the foundation model organizes samples along continuous multiomic manifolds that reflect both mutational status and downstream transcriptional programs.

### 4.3 Implications and Limitations

Methodologically, our results argue for the value of explicitly multiomic foundation models in cancer genomics. CpG-island grouping provides a scalable and interpretable representation of methylation [24, 25, 29, 30], while joint fusion with RNA-seq leverages the transcriptomic information known to be highly informative for mutation status and pathway activity [6, 19]. The fact that simple linear and shallow non-linear heads on top of frozen embeddings achieve strong performance for both cancer-type classification and mutation prediction suggests that much of the relevant structure is already captured in the foundation representation, and that task-specific re-training of the encoder may often be unnecessary.

Biologically, the joint embedding highlights both opportunities and constraints for mutation inference. For some genes and cancers, the combination of CpG-island methylation and RNA-seq yields robust zero-shot predictions, consistent with well-established epigenetic and transcriptional signatures (e.g. IDH1-associated G-CIMP in glioma, or TP53-associated changes) [18, 21, 19]. For others, performance remains modest, reinforcing the notion that not all driver alterations leave strong or consistent marks in either methylation or expression. This heterogeneity suggests that clinical applications should focus on subsets of genes and pathways with reproducible multiomic fingerprints, and that additional data types (e.g. copy number, proteomics) may be required for more comprehensive mutation inference [10, 11, 36].

Our study has several limitations. First, while we pre-train on a large multi-cohort compendium (TCGA, CPTAC-3, HCMI, TARGET), we do not explicitly construct institution- or cohort-held-out benchmarks; instead, our zero-shot evaluations rely on frozen embeddings and lightweight heads within this combined dataset. Second, although we demonstrate robustness to missing modalities at the feature and embedding level, we do not systematically dissect the marginal contribution of each modality through ablation (methylation-only vs. RNA-seq–only vs. fully multiomic). Third, interpretability of deep foundation models remains an open challenge: CpG-island grouping and pathway projections improve biological plausibility, but more targeted attribution analyses will be needed to link specific islands, genes, and pathways to model decisions. Finally, clinical translation would require careful calibration, prospective validation, and benchmarking against established molecular diagnostic methods, including targeted sequencing and panel-based tests [7, 8].

### 4.4 Future Directions

Future work can extend this framework along three main axes. First, more stringent generalization tests, including cohort-held-out and institution-held-out evaluations, as well as applications to additional public and private datasets, will be important to quantify robustness in real-world settings. Second, systematic ablations that compare methylation-only, RNA-seq only, and fully multiomic foundation models will clarify when each modality is most valuable and how their combination should be prioritized given practical constraints on data collection.

Third, integrating additional omics layers such as proteomics, copy number, and chromatin accessibility into the same foundation model could yield richer, unified representations for a broader set of downstream tasks [10, 11, 15, 36]. From a modeling perspective, extending methylation-focused foundation architectures such as *cfMethylPre*, MethylGPT, and CpGPT [33, 34, 35] to explicitly incorporate CpG-island grouping and joint multiomic supervision is a promising direction, especially when combined with improved calibration and uncertainty quantification.

Finally, the multiomic embedding learned here can be leveraged beyond mutation and lineage prediction, for example, in tumor subtype classification, prognosis, and treatment-response modeling [37, 38, 36]. In that sense, our work provides a blueprint: CpG-island grouping for scalable methylation representation, joint fusion with RNA-seq in a foundation model, and zero-shot evaluation of downstream tasks from the resulting multiomic embedding space.

## 5 Methods

### 5.1 Datasets and Availability

We utilize a multi-cohort summary of publicly available datasets spanning both training and external validation cohorts. We use all projects with paired RNA-seq and DNA methylation from TCGA, TARGET, CGCI (HTMCP-CC), the HCMI Cell Model Data Catalog (CMDC), and CPTAC-3. This includes large cohorts such as TCGA-BRCA, TCGA-KIRC, TCGA-LUAD, TCGA-UCEC and TARGET-AML, as well as rarer entities such as TCGA-ACC, TCGA-UVM, TARGET-OS, and TARGET-CCSK. For each disease where an external cohort is available (e.g. CPTAC-3 or HCMI models). From the GDC portal and associated resources, we select *all* available biospecimens with the required molecular profiles for each project, regardless of sample type (primary tumor, recurrent tumor, metastatic lesion, or normal/solid tissue controls), and handle sample-type information downstream in the analysis.

The datasets include only entries where the specific cancer is for Illumina 450K and Illumina Epic V2, taking the 450K as a base and adding the Epic V2 [39] [40]. The size of each cohort used is shown in Table 1. We tried to utilize all publicly available datasets from the GDC portal that contain either DNA methylation or RNA-seq.

**Table 1:**
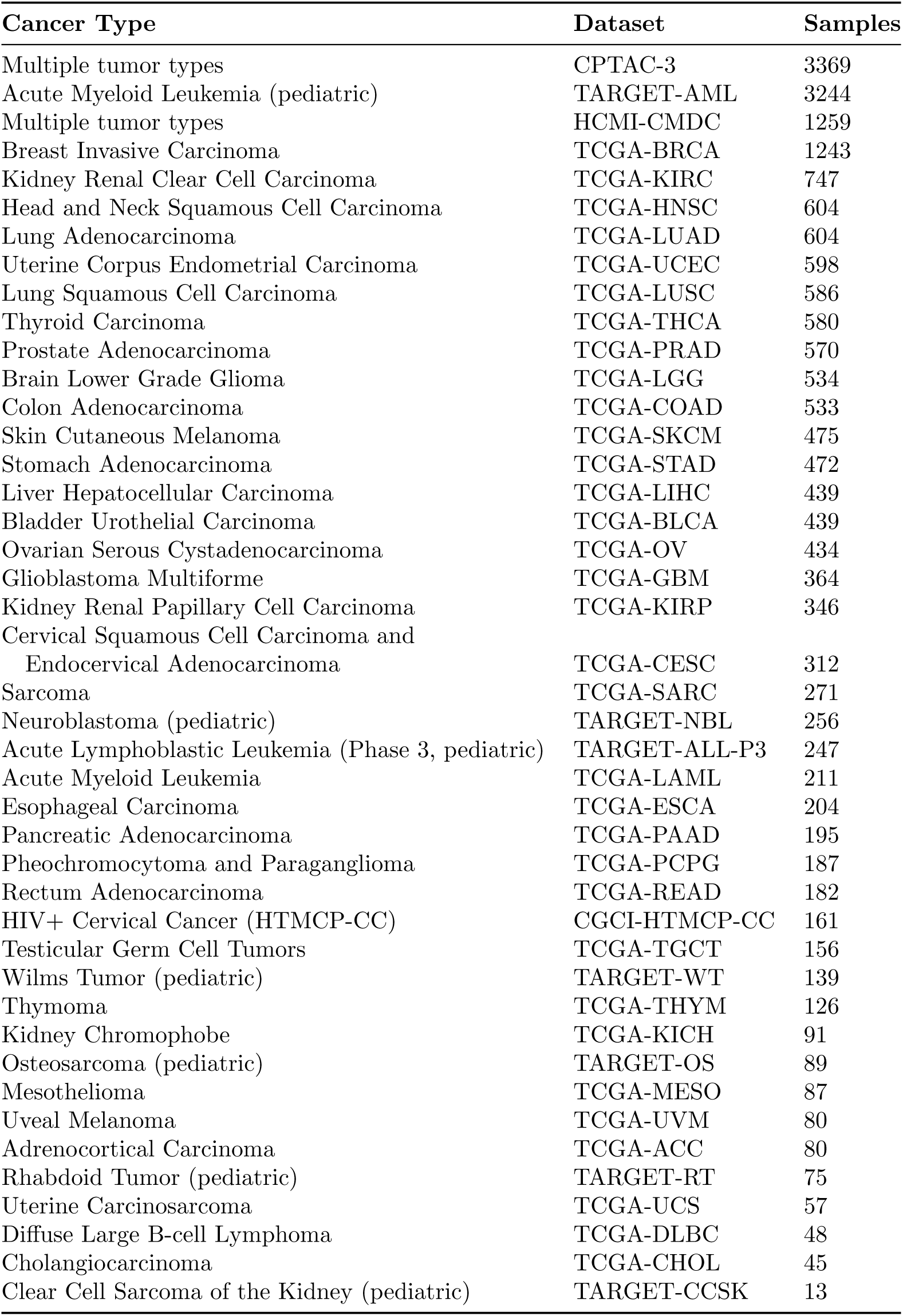
Training datasets used for foundation model construction, with sample counts.

After downloading all the cohorts, we grouped them based on cancer type. For example, CPTAC-3 contains various cancer types such as Glioblastoma Multiforme (GBM), Kung Adeonocarcinoma (LUAD) etc, which were gorouped with their corresponding TCGA, HCMI, etc. The grouping ended with samples shown in Table 2

**Table 2:** Sample counts per cohort used in the foundation model (merged multi-cohort dataset).

| Code | Full Name | Samples |
| --- | --- | --- |
| LAML | Acute Myeloid Leukemia (TCGA + TARGET) | 3854 |
| BRCA | Breast Invasive Carcinoma | 1326 |
| KIRC | Kidney Renal Clear Cell Carcinoma | 1278 |
| LUAD | Lung Adenocarcinoma | 1264 |
| UCEC | Uterine Corpus Endometrial Carcinoma | 1080 |
| CPTAC-3 | Clinical Proteomic Tumor Analysis Consortium 3 | 827 |
| COAD | Colon Adenocarcinoma | 790 |
| GBM | Glioblastoma Multiforme | 758 |
| STAD | Stomach Adenocarcinoma | 699 |
| PAAD | Pancreatic Adenocarcinoma | 642 |
| HNSC | Head and Neck Squamous Cell Carcinoma | 604 |
| LUSC | Lung Squamous Cell Carcinoma | 586 |
| THCA | Thyroid Carcinoma | 580 |
| PRAD | Prostate Adenocarcinoma | 570 |
| SKCM | Skin Cutaneous Melanoma | 553 |
| LGG | Brain Lower Grade Glioma | 534 |
| LIHC | Liver Hepatocellular Carcinoma | 439 |
| BLCA | Bladder Urothelial Carcinoma | 439 |
| OV | Ovarian Serous Cystadenocarcinoma | 434 |
| ESCA | Esophageal Carcinoma | 350 |
| KIRP | Kidney Renal Papillary Cell Carcinoma | 346 |
| CESC | Cervical Squamous Cell Carcinoma / Endocervical Adenocarcinoma | 312 |
| SARC | Sarcoma | 271 |
| READ | Rectum Adenocarcinoma | 267 |
| CMDC | Human Cancer Models Initiative (HCMi) Cell Model Data Catalog | 259 |
| NBL | Neuroblastoma (TARGET) | 256 |
| PCPG | Pheochromocytoma and Paraganglioma | 187 |
| HTMCP | HIV+ Tumor Molecular Characterization Project (Cervical Cancer) | 161 |
| TGCT | Testicular Germ Cell Tumors | 156 |
| WT | Wilms Tumor (TARGET) | 139 |
| THYM | Thymoma | 126 |
| KICH | Kidney Chromophobe | 91 |
| OS | Osteosarcoma (TARGET) | 89 |
| MESO | Mesothelioma | 87 |
| UVM | Uveal Melanoma | 80 |
| ACC | Adrenocortical Carcinoma | 80 |
| RT | Rhabdoid Tumor (TARGET) | 75 |
| UCS | Uterine Carcinosarcoma | 57 |
| DLBC | Diffuse Large B-Cell Lymphoma | 48 |
| CHOL | Cholangiocarcinoma | 45 |
| CCSK | Clear Cell Sarcoma of the Kidney (TARGET) | 13 |

### 5.2 CpG Island Grouping and Dimensionality Reduction

The dimensionality of raw Illumina HumanMethylation450 array data poses a substantial challenge, with hundreds of thousands of CpG probes per sample. To reduce complexity and enhance interpretability, we employed a CpG island mapping strategy. This approach aggregates CpG probes into biologically meaningful CpG islands, thereby reducing dimensionality while retaining regulatory signal.

The procedure begins with an annotation reference that links individual probe IDs to CpG island identifiers, genomic coordinates, associated genes, and relation-to-island categories (Island, Shore, Shelf, OpenSea) [41, 42]. Probes annotated as OpenSea, which are far from regulatory regions, are excluded to minimize noise. For each sample, probe-level beta values (0.0–1.0) are merged with the annotation and grouped by island identifier. Mean beta values across all probes in an island are then calculated to produce a single representative methylation feature.

Biologically, CpG islands are well-characterized regulatory regions, often proximal to promoters and enriched for functional epigenetic alterations in cancer. Averaging across probes within each island reduces technical variance, improves statistical power, and highlights coherent regulatory signals. In addition, relation-to-island categories (Island, Shore, Shelf) are preserved, maintaining fine-grained resolution of regulatory regions while excluding low-value probes.

The resulting data structure consists of sample-level matrices with island features and binary mutation labels for selected target genes. These island-level features formed the input for both individual per-cancer models and the TCGA-wide foundation model, enabling efficient computation, reduced memory demands, and enhanced cross-institutional generalization.

### 5.3 RNA-seq Preprocessing and Feature Selection

We processed GDC STAR gene-count TSVs listed in the project sample sheet [43, 23]. For each sample, all technical replicates were loaded, restricted to ENSG entries, and reduced to gene name and TPM columns. Within each replicate, duplicate gene rows were removed; across replicates, TPM values were averaged per gene [44]. Expression values were then log-transformed as log_2_(TPM+ 1) to stabilize variance and reduce the influence of extreme values [44]. To control input dimensionality, we restricted analysis to a whitelist of informative genes selected by a variance and expression (TPM) filter (i.e., genes consistently expressed above a TPM threshold and with high between-sample variance), typically limiting the feature set to roughly 10,000 genes [44].

### 5.4 Foundation Model Architecture

We develop a BERT-like multi-modal foundation model for cancer genomics that jointly encodes RNA-seq and DNA methylation data using self-supervised pre-training and supervised fine-tuning. The model is trained in two phases: (i) a self-supervised pre-training phase that learns general-purpose representations from large, partially labeled multiomics cohorts without using mutation labels, and (ii) a supervised fine-tuning phase in which a mutation-prediction head is trained (or adapted) on top of the frozen or partially unfrozen encoder.

The model operates on two input modalities:

- **RNA-seq:** gene-level expression profiles for 10,000 genes.
- **DNA methylation:** aggregated beta values for 26,086 CpG islands.

During fine-tuning, the model outputs mutation probabilities for 214 target genes in a multi-task setting (one binary output per gene).

#### 5.4.1 Modality-Specific MLP Encoders

Each modality is first processed by an asymmetric multilayer perceptron (MLP) encoder that compresses the high-dimensional input into a *d*_model_ = 768-dimensional embedding. The methylation encoder is allocated substantially more capacity than the RNA encoder, reflecting its larger input dimensionality (26k vs. 10k features).

**RNA encoder** For an RNA input vector **x**^rna^ ∈ ℝ^10,000^, the encoder applies three linear blocks with layer normalization, ReLU activation, and dropout (rate 0.1):

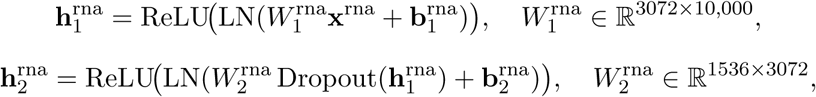

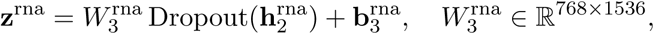

yielding a 768-dimensional RNA embedding **z**^rna^.

**Methylation encoder** For a methylation input vector **x**^meth^ ∈ ℝ^26,086^, the encoder uses a deeper and wider MLP:

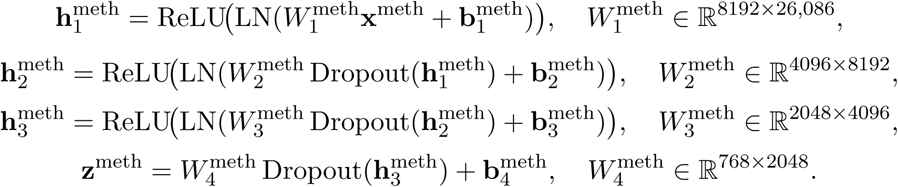

This yields a 768-dimensional methylation embedding **z**^meth^. The asymmetric design (larger hidden layers for methylation) provides roughly seven-fold more parameters in the methylation encoder, accommodating its higher input dimensionality while maintaining a shared embedding size.

#### 5.4.2 Transformer Fusion with Missing-Modality Support

The core fusion block is a transformer-based module with self-attention and cross-attention, operating on one token per modality (the 768-dimensional embeddings). To support incomplete multiomics profiles, the model includes learned missing-modality tokens and modality-aware scaling.

**Learned missing tokens** We introduce two learned vectors:

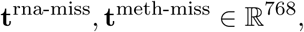

initialized with small random values (standard deviation 0.02). For a given sample, if RNA-seq (resp. methylation) is not available, **z**^rna^ (resp. **z**^meth^) is replaced by the corresponding missing token before entering the transformer. These tokens are trained jointly with the rest of the model and enable the network to handle mixed batches where some samples have both modalities and others only one.

**Modality-specific transformer encoders** Each modality is first processed by its own transformer encoder consisting of *L* = 6 layers, model dimension *d*_model_ = 768, *n*_head_ = 12 attention heads, and feed-forward dimension *d*_ff_ = 3072:

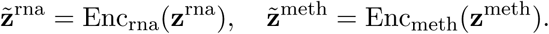

Each layer follows the standard transformer block with multi-head self-attention, residual connections, layer normalization, and a two-layer feed-forward network with ReLU activation [45].

**Cross-attention fusion** To capture cross-modal interactions, we apply bidirectional cross-attention:

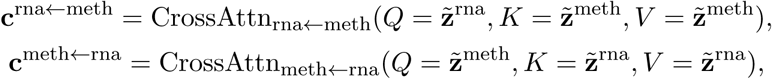

where each cross-attention is implemented as a multi-head attention layer with 12 heads. Residual connections and layer normalization yield the fused representations:

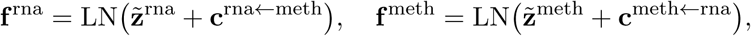

where **f** ^rna^, **f** ^meth^ ∈ ℝ^768^ are modality-specific embeddings enriched with cross-modal context.

#### 5.4.3 Self-Supervised Pre-Training Heads

Pre-training uses four self-supervised objectives: two masked-value reconstruction tasks and two cross-modal prediction tasks, in the spirit of masked language modeling [46] and recent omics foundation models [47, 48].

**Masked value reconstruction** For each sample, we randomly mask 15% of RNA and methylation features (set to zero or a mask embedding at the input) and ask the model to reconstruct the original values.

From **f** ^rna^, an MLP of the form

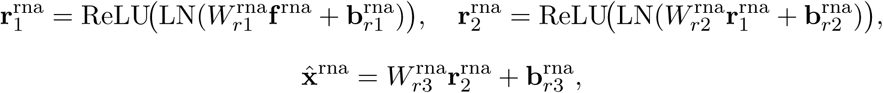

maps 768 → 2048 → 1024 → 10,000; an analogous head is used for methylation (768 → 2048 → 1024 → 26,086). Losses are mean squared error (MSE) computed only on the masked positions:

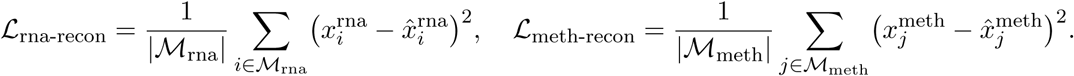

**Cross-modal prediction** To encourage the model to learn cross-modal dependencies, we add two auxiliary heads:

- RNA-from-methylation: predicts full RNA expression **x̃**^rna^ from **f** ^meth^.
- Methylation-from-RNA: predicts full methylation profile **x̃**^meth^ from **f** ^rna^.

Both heads use the same 768–2048–1024–output architecture and MSE loss over all observed features:

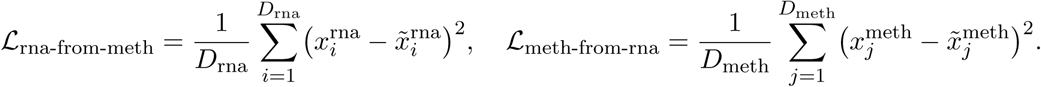

**Pre-training loss** The overall pre-training loss combines these four objectives:

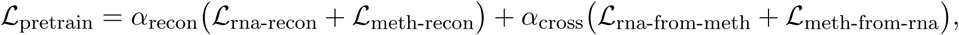

with default weights *α*_recon_ = 1.0 and *α*_cross_ = 0.5. All encoder, transformer, missing-token, and pre-training head parameters are updated during this phase.

#### 5.4.4 Mutation Prediction Head and Fine-Tuning

For supervised mutation prediction, we concatenate the fused embeddings from both modalities:

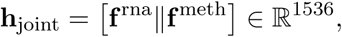

and feed this through a two-layer MLP:

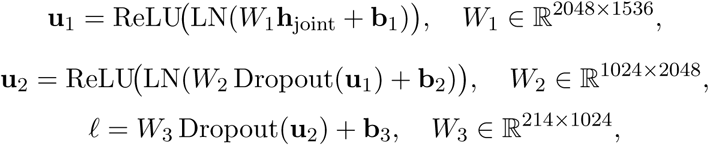

producing logits *ℓ_g_* for each gene *g* = 1*, . . .,* 214.

**Modality-aware scaling** To adjust prediction confidence based on which modalities are present, we apply a learned scalar scaling to the logits:

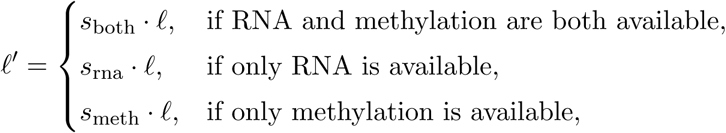

where *s*_both_*, s*_rna_*, s*_meth_ are learned parameters. Mutation probabilities are then

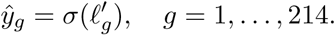

We optimize a binary cross-entropy loss with logits in a multi-task, multi-label setting, with per-gene class weights derived from mutation prevalence and capped to the range [1.0, 10.0] to mitigate imbalance.

**Training regimes** We consider two fine-tuning regimes:

- **Frozen encoder** (default): the RNA and methylation MLP encoders and transformer fusion module are frozen at their pre-trained weights, and only the mutation head (and modality scaling factors) are trained. This reduces the number of trainable parameters, mitigates overfitting on small cohorts, and preserves pre-trained representations.
- **End-to-end fine-tuning**: all layers, including encoders and transformers, are unfrozen and trained with a smaller learning rate. This can yield improved performance when sufficient labeled data are available.

In both cases, we use the Adam optimizer with weight decay, dropout of 0.1 throughout, early stopping based on validation loss, and three random seeds to report mean ± standard deviation of evaluation metrics.

### 5.5 Cancer Type Classification from Frozen Foundation Embeddings

To assess whether the pretrained foundation model captures cancer-type information, we trained a linear probe on frozen embeddings to perform cancer-type classification. Linear probes, which are simple classifiers that fit on top of frozen representations, are a standard tool to quantify the quality of learned embeddings in vision and language models [49, 50] and have been increasingly adopted in biological foundation models [31].

#### 5.5.1 Data and Embedding Extraction

We used the unpaired foundation data dataset and retained samples with non-missing cancer type labels, excluding rows labeled as Unknown. Samples with both rna available = False and meth available = False were removed.

Feature selection matched the main experiments:

- **RNA features:** non-chr* columns after excluding metadata columns (e.g. sample identifiers, file-related columns, availability flags, and the cancer type label), and discarding columns with non-gene naming conventions (e.g. lowercase).
- **Methylation features:** columns with names beginning with chr, corresponding to CpG-island methylation features.

All features were cast to float32 and cleaned using nan-to-zero replacement for numerical stability. The number of columns was trimmed to match the model’s expected rna input dim and meth input dim.

We loaded the best pretrained checkpoint (with the mutation head disabled or set to num genes for mutation = 0) and used the model’s encoding interface to compute fused em-beddings:

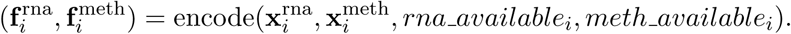

Unavailable modalities were handled via the learned missing-modality tokens as in the main architecture. The final representation for each sample was the concatenation

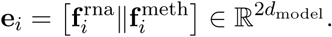

Embeddings were sanitized using np.nan to num prior to downstream analysis.

#### 5.5.2 Linear Probe and Cross-Validation

On top of the frozen embeddings, we trained a multinomial logistic regression classifier (linear probe) for cancer-type prediction [49, 50]. Cancer types were encoded as integer-class labels. Within each cross-validation fold, embeddings were standardized (zero mean, unit variance) using statistics computed on the training split and applied to the validation split.

We used stratified *K*-fold cross-validation (default *K* = 5). If any cancer type had fewer than *K* samples, *K* was reduced accordingly to ensure each class appeared in every fold. For each fold, the classifier was trained on the training split and evaluated on the held-out fold. The logistic regression probe was implemented with:

- multinomial loss with softmax,
- *ℓ*_2_ regularization with *C* = 1.0,
- class-weighting set to balanced to compensate for imbalanced cancer-type frequencies,
- maximum 1000 iterations.

#### 5.5.3 Evaluation Metrics and Visualization

For each fold, we computed:

- overall accuracy and balanced accuracy,
- macro- and weighted-averaged F1 scores,
- macro-averaged one-vs-rest AUC using predicted class probabilities,
- per-class precision, recall, F1, and support.

Metrics were aggregated across folds to report mean ± standard deviation per metric. We further visualized misclassification structure using confusion matrices (both raw counts and row-normalized).

To visualize how the foundation embeddings organize cancer types, we applied principal component analysis (PCA) [51] to the concatenated embeddings and generated two-dimensional scatterplots (PC1 vs. PC2) colored by cancer type. Clear clustering by cancer type in this space indicates that the pretrained encoder captures disease-specific structure even without supervision from mutation labels.

### 5.6 Pathway Enrichment Analysis on Foundation Embeddings

We next investigated whether the foundation embeddings correlate with known biological pathways, focusing on cancer-relevant gene sets. We used the Hallmark gene sets from the Molecular Signatures Database (MSigDB) [52] and, when indicated, additional KEGG [53] and Gene Ontology (GO) biological-process gene sets [54]. Pathway activity was estimated using a single-sample gene set enrichment strategy similar to ssGSEA [55].

#### 5.6.1 Embedding Extraction and Pathway Scoring

We used the same multiomics dataset as in the cancer-type classification analysis, retaining samples with at least one modality available (RNA or methylation). RNA and methylation features were selected and preprocessed as described above, and fused embeddings

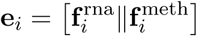

were extracted from the frozen pretrained model.

For pathway scoring, we restricted to genes present in both the expression matrix and the chosen gene sets. Gene-level expression was first z-scored across samples. For each pathway *p* with gene set *G_p_*, and each sample *i*, we defined a simple single-sample pathway activity score:

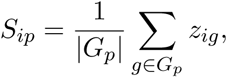

that is, the mean z-scored expression of member genes in that sample. This yields a pathway activity matrix *S* ∈ ℝ*^N^*^samples×^*^N^*^pathways^ .

#### 5.6.2 Embedding–Pathway Correlation

To relate the learned embeddings to pathway activity, we computed the Pearson correlation between each embedding dimension and each pathway score across samples, skipping any dimension or pathway with zero variance. Let **e**_·_*_d_* denote embedding dimension *d* across samples and *S*_·_*_p_* the activity of pathway *p*; the correlation matrix

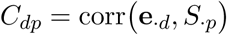

summarizes the association between embedding axes and pathways.

We used *C* to:

- identify embedding dimensions with the highest mean absolute correlation across pathways,
- identify pathways with the highest mean absolute correlation across embedding dimensions,
- visualize the top-dimensional submatrix of *C* as a clustered heatmap.

These analyses highlight embedding directions that align with coherent biological processes (e.g. proliferation, immune response, hypoxia).

#### 5.6.3 Pathway Co-activation and Cancer-Specific Activity

We examined pathway co-activation patterns by computing pairwise Pearson correlations among pathway activity scores across samples, focusing on the most variable pathways. The resulting pathway–pathway correlation matrix was visualized as a heatmap to identify modules of co-regulated pathways.

To study pathway differences between cancer types, we aggregated pathway scores by cancer type and computed mean

## 6 Conclusion

We presented a pan-cancer, multiomic foundation model that jointly encodes CpG-island DNA methylation and RNA-seq profiles into a shared latent space. By aggregating probe-level methylation into biologically meaningful CpG islands and reducing RNA-seq to a curated set of high-variance genes, we achieved substantial dimensionality reduction while retaining regulatory signal suitable for large-scale representation learning. A BERT-like transformer with modality-specific encoders, self-supervised reconstruction and cross-modal objectives, and explicit support for missing modalities yields compact embeddings that integrate epigenetic and transcriptomic information across diverse cancer types.

Zero-shot evaluations demonstrate that these embeddings support accurate cancer-type classification via a linear probe and strong mutation prediction for many gene–cancer pairs using shallow classifiers, without task-specific re-training of the encoder. Embedding visualizations and pathway analyses further show that hallmark oncogenic and immune programs are organized as smooth gradients in the latent space, indicating that the model captures meaningful biological structure rather than merely fitting labels.

Together, these results highlight the value of combining CpG-island grouping with multiomic foundation modeling for mutation and lineage inference. Our framework provides a scalable starting point for building more comprehensive multiomic foundation models, and for extending their use to downstream tasks such as subtype classification, prognosis, and treatment-response prediction in precision oncology.

